# Bent Hockey-Stick Stock-Recruitment Function

**DOI:** 10.64898/2025.12.17.695007

**Authors:** Hiroshi Okamura, Shota Nishijima, Momoko Ichinokawa

## Abstract

The hockey-stick is a simple and widely applicable stock–recruitment function. One drawback is its non-differentiability at the breakpoint, which can be especially problematic when using state-space models for stock assessment. Here, we develop a new bent hockey-stick model that retains the desirable properties of the original hockey-stick function while providing a differentiable breakpoint. We also propose a new algorithm for estimating MSY under multiplicative process stochasticity in the stock–recruitment relationship. Simulation studies demonstrate the utility of these approaches, which can substantially enhance the robustness and sustainability of fisheries management.

## Introduction

Estimating the stock-recruitment (SR) relationship is central to sustainable fisheries management (Subbey et al., 2014). However, parameter estimation in SR models is notoriously difficult (Quinn and Deriso, 1999; Walters and Martell, 2003). Because fish recruitment is strongly influenced by environmental variation, SR data typically exhibit large fluctuations and shotgun-like scatterplots. When estimating SR relationship, inaccurate and imprecise parameter estimates – and consequently, unreliable biological reference points – often arise from statistical issues such as errors-in-variables and time-series bias. As a result, regardless of the true underlying SR relationship, the estimated SR curve often appears flat or linearly increasing within the observed range of spawning stock biomass (SSB), particularly when conventional models such as the Beverton–Holt or Ricker functions are used (Ichinokawa et al., 2017; Okamura et al., 2017). With a flat SR curve, recruitment remains constant, so increasing the fishing rate does not reduce future recruitment unless the stock is driven to collapse. In contrast, with a linearly increasing SR curve, recruitment is expected to level off only far above the observed range of SSB, which leads to reducing the fishing rate to rebuild the stock toward such high levels. Consequently, assuming a flat SR relationship can lead to overly aggressive fishing rates, whereas assuming a linear SR relationship can lead to overly conservative management.

The Hockey-Stick (HS) SR relationship, a simple segmented regression model, can circumvent some of the problems inherent in the Beverton-Holt (BH) and Ricker (RI) models (Ichinokawa et al., 2017). When the flat SR relationship is indicated, the HS model can set the breakpoint at the minimum observed SSB, which leads to a conservative management outcome even if the true breakpoint lies below this value. Conversely, when a linearly increasing SR relationship is indicated, the HS model can set the breakpoint at the maximum observed SSB. This prevents fishery closures and enables adaptive management by allowing SSB to increase through reduced fishing pressure until the true breakpoint becomes apparent.

However, the HS model has a drawback of being non-differentiable at the breakpoint (Barrowman and Myers, 2000; Mesnil and Rochet, 2010). This issue becomes more critical when a state-space model is used for stock assessment (e.g., Nielsen and Berg, 2014; Nishijima et al., 2021), because the breakpoint must be estimated for latent variables, while many optimization and Bayesian estimation frameworks assume differentiability with respect to parameters (Kristensen et al., 2016; Carpenter et al., 2017). Mesnil and Rochet (2010) proposed a bent-hyperbola model as a differentiable HS-like SR function, hereafter referred to as MR. Although this approach is both practical and appealing, it does not strictly preserve the HS property; recruitment becomes only approximately flat as SSB increases, and becomes completely flat only in the limit as SSB approaches infinity. If a differentiable approximation to the HS model that remains exactly horizontal as SSB increases could be developed, it would be more suitable for application within a state-space framework and would yield more stable biological reference points.

We propose a bent HS (BHS) function that is differentiable at the breakpoint and approaches a plateau for large SSB values. The performance of the BHS model is evaluated by applying a state-space model with the BHS-SR function to simulated data based on the Falkland Islands’ *Loligo gahi* squid stock assessment (McAllister et al., 2004). Estimation of biological reference points related to maximum sustainable yield (MSY) is generally known to be sensitive to process noise in the SR model (Bousquet et al., 2008; Bordet and Rivest, 2014). To address this, we develop a simple numerical optimization method for estimating biological reference points under stochastic environments. For comparison, the HS, MR, BH, and RI models are also applied to the same simulated dataset. Finally, the estimated biological reference points are compared across all models.

## Materials and Methods

### A Bent Hockey-Stick Stock-Recruitment Function

We introduce a BHS-SR function, defined by

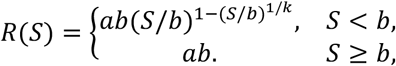

where *R* is recruitment, *S* is SSB, *a* is the maximum population growth rate, *b* is the breakpoint corresponding to the threshold spawning biomass above which recruitment becomes constant, and *k* is a smoothing parameter that connects the curve for *S* < *b* with the constant *ab* for *S* ≥ *b*. When *k* → 0, the model reduces to the ordinary HS-SR function (Table 1), whereas *k* → ∞ results in constant recruitment.

**Table 1.** Estimated parameters and corresponding biological reference points derived from the stock–recruitment (SR) relationships used in this study. HS: Hockey-Stick; MR: Mesnil-Rochet; BHS: Bent Hockey-Stick; BH: Beverton-Holt; RI: Ricker.

| Model | Equation for $R$ | $a$ | $b$ | $\sigma$ | $F_{MSY}$ | $S_{MSY}$ |
| --- | --- | --- | --- | --- | --- | --- |
| HS | $\begin{cases} aS & (S < b), \\ ab & (S \geq b) \end{cases}$ | 192.0 | 20.1 | 0.71 | 0.87 | 44.1 |
| MR | $a \left[ S + \sqrt{b^2 + \gamma^2/4} - \sqrt{(S - b)^2 + \gamma^2/4} \right]$ | Not Converged | | | | |
| BHS | $\begin{cases} \frac{1}{k_m} \int_0^{k_m} ab \left( \frac{S}{b} \right)^{1 - \left( \frac{S}{b} \right)^{\frac{1}{z}}} dz & (S < b), \\ ab & (S \geq b) \end{cases}$ | 98.2 | 39.4 | 0.71 | 0.86 | 44.8 |
| BH | $\frac{aS}{1 + S/b}$ | 476.7 | 8.6 | 0.72 | 0.84 | 48.3 |
| RI | $aS \exp(-S/b)$ | 179.7 | 113.4 | 0.17 | 0.85 | 95.2 |

When one differentiates the ordinary HSSR with respect to *b*, the left derivative is zero and the right derivative is *a*. In other words, ∂*R*/∂*b* = 0 when *S* < *b*, whereas ∂*R*/∂*b* = *a* when *S* ≥ *b*. Because the left and right derivatives are unequal, the ordinary HSSR is non-differentiable with respect to *b*.

By contrast, for the BHS function, differentiating *R* with respect to *b* for *S* < *b* yields

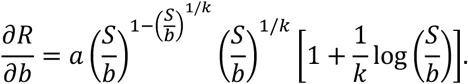

When *S* approaches *b*, the second term in the square bracket tends to zero. Hence, ∂*R*/∂*b* = *a* holds on both sides of *S* = *b*.

When we introduce the parameter *k* into the SR function, the number of parameters to be estimated increases to three. However, including a shape parameter to form a three-parameter stock–recruitment relationship often causes identifiability and convergence problems, because typical stock–recruitment data rarely contain sufficient information to estimate all three parameters reliably (Brooks et al., 2019). Therefore, a two-parameter SR function is generally preferable. To account for the unknown *k*, we instead use an averaged recruitment function for *S* < *b*, assuming a uniform prior distribution for *k*:

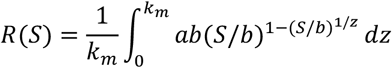

where we temporarily set *k*_*m*_ = 5. Integrating over *k*_*m*_ stabilizes parameter estimation even when the available time series is short. As *k*_*m*_ increases, the left side of the curve (below the breakpoint) becomes more rounded (Fig. 1). Preliminary analyses of identifiability and sensitivity of the *k*_*m*_ parameter, as well as practical approaches for its selection, are summarized in the Supplementary Online Materials.

**Figure 1.**
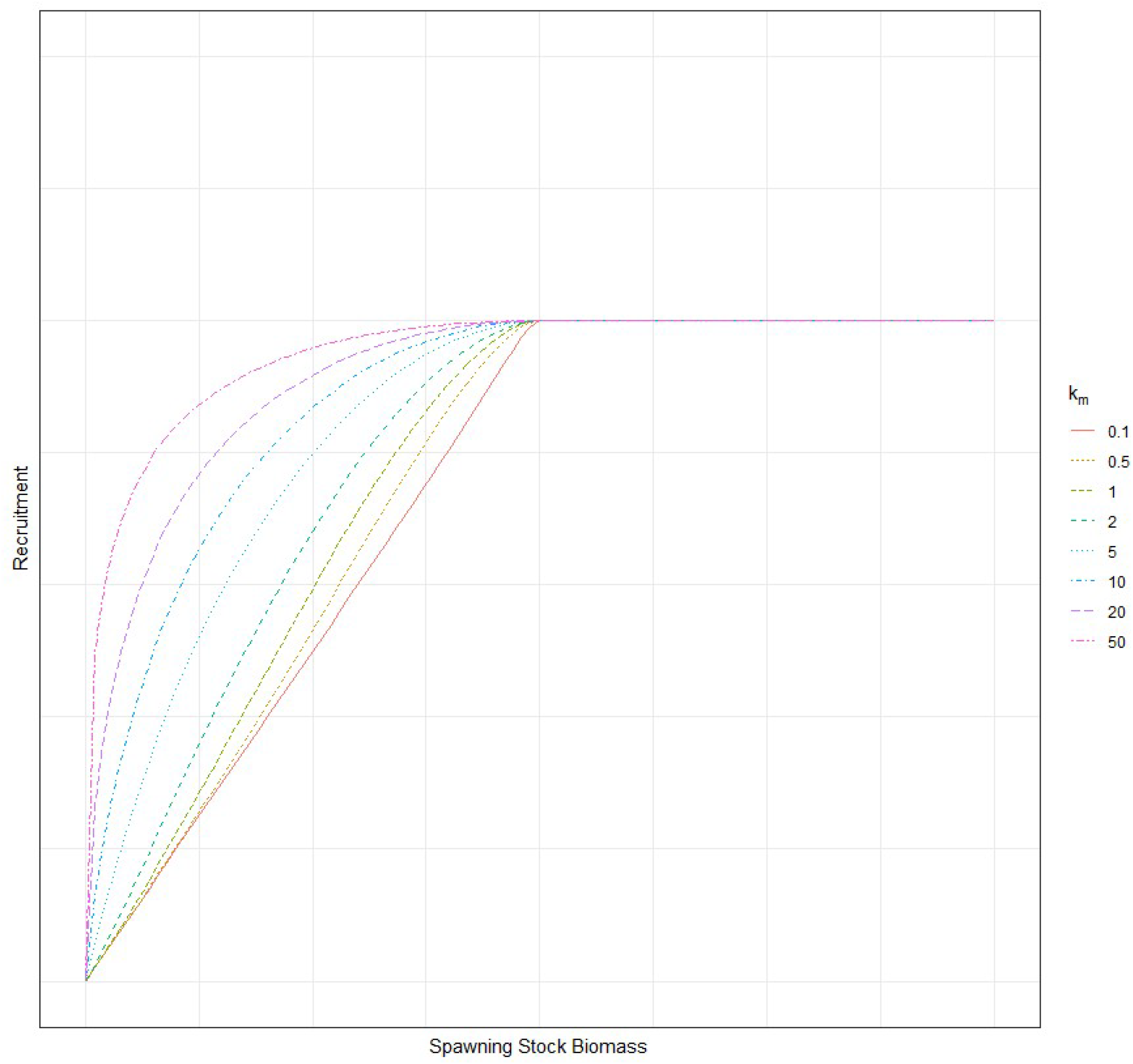
Shapes of bent hockey-stick curves for various *k*_*m*_ values.

### A state-space Leslie depletion model for the Falkland Islands’ Loligo gahi squid

We use total catch and catch-per-unit-effort (CPUE) data for the Falkland Islands’ *Loligo gahi* squid from 1987 to 2000 (Table 1 in McAllister et al., 2004). Because the dataset includes a time series of average individual weight, we convert biomass-based quantities to abundance-based quantities by dividing both CPUE and catch by the corresponding average weight.

The within-year population size (*N*) is updated by accounting for natural mortality and fishing removals (*U*) in each weekly interval (Δ*w*):

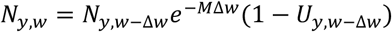

where *y* denotes year, *w* denotes week, and *M* is the instantaneous natural mortality rate per week, set to 7×0.0133 according to Roa-Ureta and Arkhipkin (2007). This model corresponds to a so-called Leslie model (Leslie and Davis, 1939). The initial population size with a density-dependent stock-recruitment relationship is modelled as

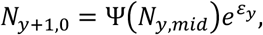

where Ψ is a stock-recruitment function, which can take the form of HS, MR, BHS, BH, or RI. *N*_*y,mid*_ = *N*_*y*,0_*e*^‒*MT*^ ∏(1 − *U*_*y,w*_) represents the population size at the middle of the year (*T* = 26), after the end of the fishing season, which serves as the spawning stock in our model. Here *U*_*y, w*_ = *C*_*y, w*_ /*N*_*y, w*_ (where *C*_*y, w*_ is the catch at week *w* in year *y*), and ε_y_∼*Normal*(0, σ^2^). Taking the natural logarithm of both sides of the equation,

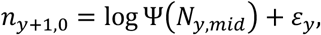

where *n*_*y, w*_ = log (*N*_*y, w*_). For the initial year (*y* = 0), we assume *n*_0,0_∼*Normal*(*ñ*_0_, σ^2^).

We assume that CPUE is proportional to population size with lognormal observation error:

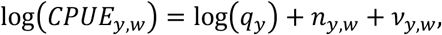

where ν_*y, w*_ ∼*Normal*(0, τ^2^).

The logarithm of fishing efficiency *q*_*y*_ is assumed to follow a random walk:

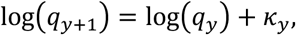

with 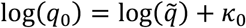, and *κ*_*y*_∼*Normal*(0, η^2^).

Because the log population size *n*_*y,0*_ and the log fishing efficiency log(*q*_*y*_) are random effects (unobserved latent variables), they must be integrated out when maximizing the marginal likelihood. We use an R package TMB (Template Model Builder: Kristensen et al., 2016) to maximize the marginal likelihood via the Laplace approximation.

### Estimation of biological reference points under a stochastic environment

Because the BHS has a complex nonlinear functional form, it is difficult to obtain closed-form solutions for MSY-related biological reference points. Therefore, numerical approximation methods must be used to calculate these quantities.

To estimate the equilibrium population size 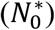 under a constant fishing rate (*F*) in a stochastic environment, we need to solve the following equation:

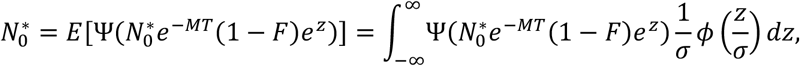

where *ϕ* denotes the probability density function of the standard normal distribution, and *σ* is the standard deviation of the random variable *z*. Note that *N*_0_^*\**^ is a function of the constant annual fishing rate *F*. The constant annual fishing rate *F* is defined as *F* = 1 − (1 − *U*)^*T*^, where *U* is a constant weekly fishing rate. Because this equation is nonlinear and has no closed-form solution, we use numerical optimization to solve for *N*_0_ ^*^. The objective function is defined on a log-scale; that is, we minimize

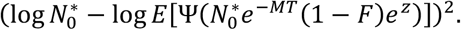

We use an *n*-point Gauss-Hermite quadrature to approximate the above expectation:

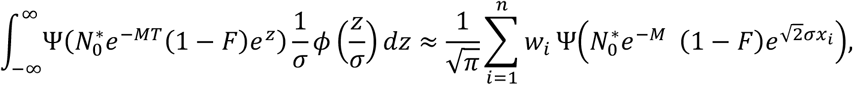

where the nodes *x*_*i*_ are the roots of the Hermite polynomial and *w*_*i*_ are the corresponding weights.

The fishing rate at MSY is estimated by maximizing 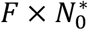:

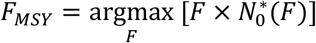

where we explicitly denote *N*_0_^*^ as a function of *F* indicating that the equilibrium population size depends on *F*.

This is generally a complex nonlinear equation, and therefore we use a numerical optimization procedure again. Because we need numerical optimization to solve for *N*_0_ ^*^ as mentioned above, a nested numerical optimization is required to solve for *F*_*MSY*_. Given *F*_*MSY*_, other biological reference points are calculated as follows:

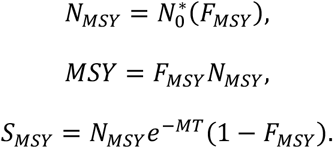

We hereafter refer to this new procedure as the optimization-based stochastic MSY estimation algorithm. To evaluate the appropriateness of the above calculation, the optimization-based stochastic surplus productions 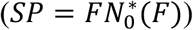 are compared with the surplus productions obtained from *G* = 3,000 simulation runs. Parameter estimates are taken from the Falkland Islands’ *Loligo gahi* squid assessment based on various SR relationships.

In the simulation, the log initial population size for each year *y* and simulation *g* is updated as

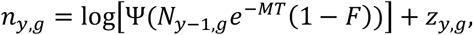

where *n*_*y,g*_ = log *N*_*y,g*_ and *z*_*y,g*_∼*N*(0, *σ*^2^). The simulation-based stochastic surplus production is calculated as the ensemble mean over simulations,

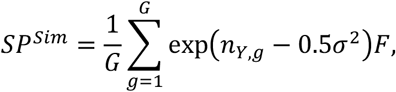

for *Y* = 100 (i.e., the 100th year of simulation). Because a log-scaled objective function is used in our algorithm, the simulated biological reference points are adjusted using a mean correction (Bousquet et al., 2008; Thorson and Kristensen, 2016) to account for the log-normal distribution.

To investigate the impact of incorporating stochasticity on the estimation of biological reference points, we also calculate the optimization-based deterministic surplus production by setting the process standard deviation (*σ*) to zero in the above equation.

To facilitate the application of the BHS model and the new optimization-based stochastic MSY estimation algorithm and to help reproduce our results, the program code is publicly available on the first author’s website (https://github.com/OkamuraHiroshi/Bent-Hockey-Stick-Stock-Recruitment-Model).

### A simulation test

The performance of the BHS-SR relationship is evaluated using simulated data. In the simulations, one of four SR models – the HS, BHS, BH, and RI – is assumed to represent the true SR relationship (Table 1; the MR model is excluded because it fails to converge for the original dataset). The parameters of the assumed SR model are estimated by fitting the state-space model (SSM) with that SR function to the Falkland Islands’ *Loligo gahi* squid data.

For the simulated datasets, sets of parameters are first generated from a multivariate normal distribution with the estimated log-parameters and their variance-covariance matrix. The population size in each year is then simulated by removing the proportion caught by fisheries. Logit-transformed fishing rates are randomly drawn from a normal distribution based on the estimated logit-transformed fishing rates from the original dataset, and then inverse-logit transformed.

Some of the simulated population trajectories are implausible because random draws of parameters and fishing rates may yield unrealistically high or low population sizes. To exclude such unrealistic trajectories, we resample the simulated population time series with the weight

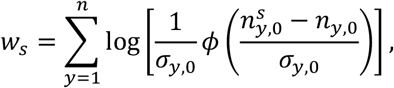

where *n*_*y*,0_ = log *N*_*y*,0_ is the log of the population size at the start of the fishing season in year *y*, obtained from model fitting to the original dataset; σ_*y*,0_ is the corresponding standard error of *n*_*y*,0_; and 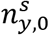 is the log of the *s*-th simulated population size at the start of the fishing season in year *y*. This resampling procedure effectively conditions the simulated time series on the original data and produces plausible population trajectories that remain close to the original one. We first generate 1,000 unconditioned simulated datasets, and then resample 100 conditioned datasets without replacement.

After generating the plausible population trajectories, the observed CPUEs proportional to the population size are randomly sampled using the parameter estimates from the original dataset. The simulated catch and CPUE time series are then fitted to the model assuming the HS, MR, BHS, BH, or RI SR relationship. The parameters used in the simulation are employed as initial values. Convergence is evaluated based on whether (1) the change in log-likelihood is sufficiently small (< 10^-10^%), (2) the maximum gradient is sufficiently small (< 0.01), (3) the Hessian matrix is positive definite, and (4) all elements of the variance–covariance matrix are numeric (i.e. not NA). When convergence fails, the initial values are modified and the estimation is repeated. For the MR and BHS models, the smoothness-adjustment parameters (*γ* for MR and *k*_*m*_ for BHS) are also adjusted if the default setting fails to converge. Specifically, for *γ*, the default value is 0.01 and alternative values of 0.1 and 0.5 are tested; for *k*_*m*_, the default is 5 and values between 0.1 and 20 are examined.

Because there are four SR models assumed as the true model and five SR models used for estimation, a total of 4×5 = 20 model combinations are evaluated. The performance of each estimation model is assessed based on the relative bias of biological reference points for *F*_*MSY*_ and *S*_*MSY*_, calculated as (EST^(*s*)^ − TRUE^(*s*)^)/TRUE^(*s*)^.

## Results

The state-space stock assessment models applied to the Falkland Islands’ *Loligo gahi* squid dataset converged successfully for all SR functions except MR. The HS and BHS models produced similar SR relationships, although the breakpoint of the BHS model shifted slightly to the right compared to the HS model (Fig. 2). The estimated *F*_*MSY*_ values were similar across all models, whereas the estimated *S*_*MSY*_ value for the RI model was much larger than those for the other models (Table 1). The estimated states (pairs of *R* and *S*) were similar among the HS, BHS, and BH models, but somewhat different for the RI model (Fig. 2).

**Figure 2.**
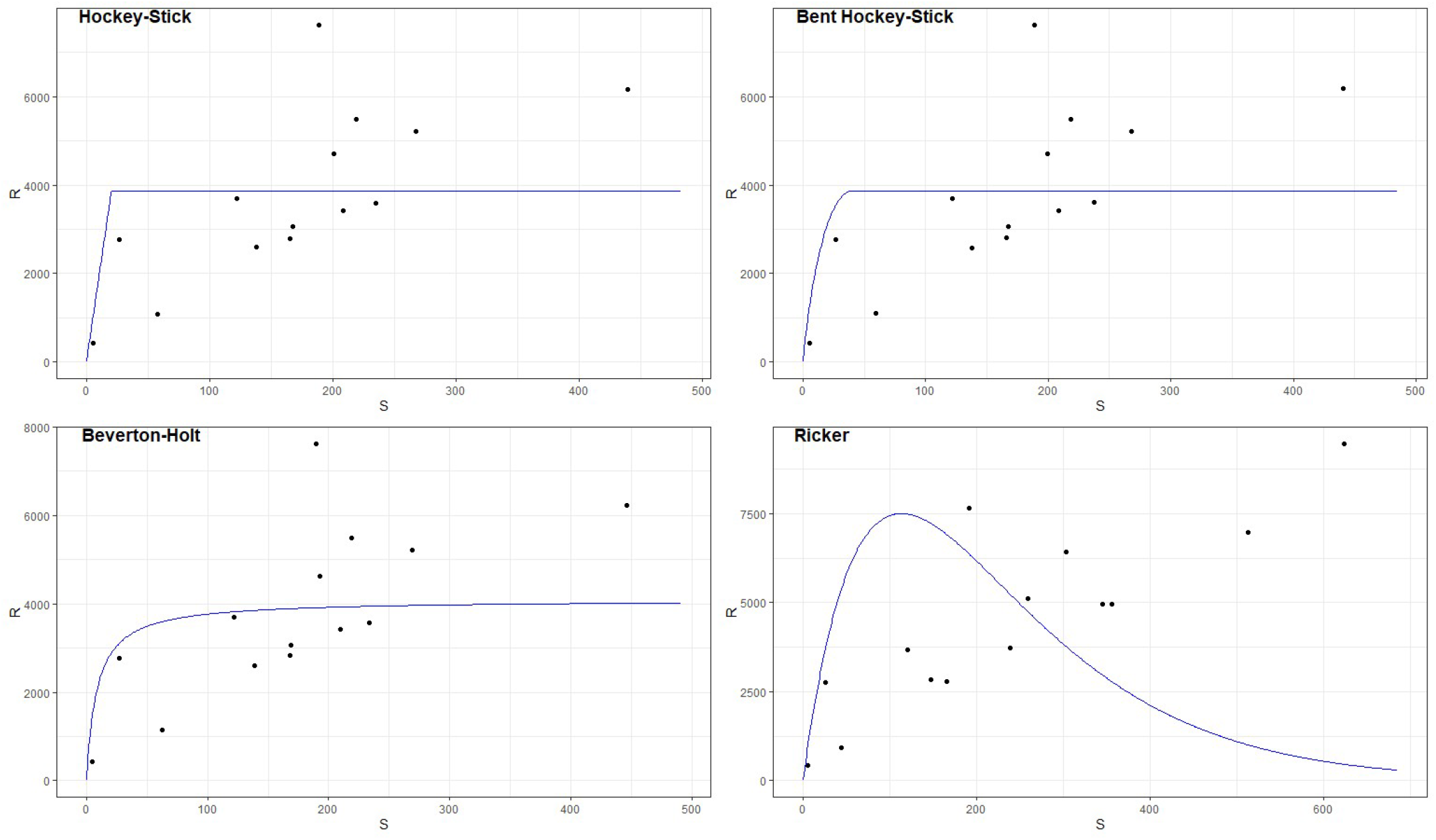
Stock–recruitment (SR) relationships estimated from the Falkland Islands’ *Loligo gahi* squid data. The solid line represents the fitted SR curve, and the points indicate spawning biomass and recruitment estimates from the state-space model.

Surplus productions curves as a function of fishing rate were compared among the stochastic simulation, optimization-based deterministic, and optimization-based stochastic calculations, all conducted using the estimated parameters. The results from the simulation-based and optimization-based stochastic calculations were very similar, whereas those from optimization-based deterministic calculation differed noticeably, particularly for the HS and BHS models (Fig. 3). When process noise was ignored in the HS and BHS models, the fishing rate at MSY (*F*_*MSY*_) tended to be overestimated, indicating that neglecting stochasticity can be more problematic than in the BH and RI models. In contrast, the BH and RI models were relatively insensitive to whether stochasticity was considered or not, although the apparent insensitivity of the RI model likely reflects its relatively small estimated process standard deviation (Table 2).

**Table 2.** Simulation results. Converge: percentage of converged results; RBF: mean relative bias (%) of fishing rate at MSY; RBS: mean relative bias (%) of spawning stock biomass at MSY. The values in parentheses indicate standard deviations of relative bias (%).

| SR function | True Model |  |  |  |  |  |  |  |  |  |  |  |
| --- | --- | --- | --- | --- | --- | --- | --- | --- | --- | --- | --- | --- |
|  | Hockey-Stick |  |  | Bent Hockey-Stick |  |  | Beverton-Holt |  |  | Ricker |  |  |
|  | Converge | RBF | RBS | Converge | RBF | RBS | Converge | RBF | RBS | Converge | RBF | RBS |
| HS | 69 | -6.7(10.6) | 161(264) | 80 | -6.9(14.7) | 131(197) | 69 | -1.7(14.4) | 81(150) | 96 | 1.5(6.2) | -23(69) |
| MR | 100 | -6.9(9.4) | 188(473) | 100 | -6.6(12.8) | 121(161) | 100 | -1.4(11.3) | 61(122) | 99 | 3.6(6.0) | -42(25) |
| BHS | 100 | -3.7(9.3) | 110(318) | 100 | -1.9(14.7) | 91(182) | 100 | 2.9(17.3) | 53(183) | 100 | -0.2(5.1) | -22(31) |
| BH | 98 | -1.3(11.7) | 55(335) | 97 | 3.7(14.2) | 19(154) | 92 | 7.8(14.6) | -7(121) | 100 | -1.2(7.0) | -7(52) |
| RI | 70 | -12.8(9.2) | 398(655) | 62 | -10.7(11.9) | 263(295) | 64 | -5.9(13.1) | 202(210) | 26 | -0.1(2.0) | 15(22) |

**Figure 3.**
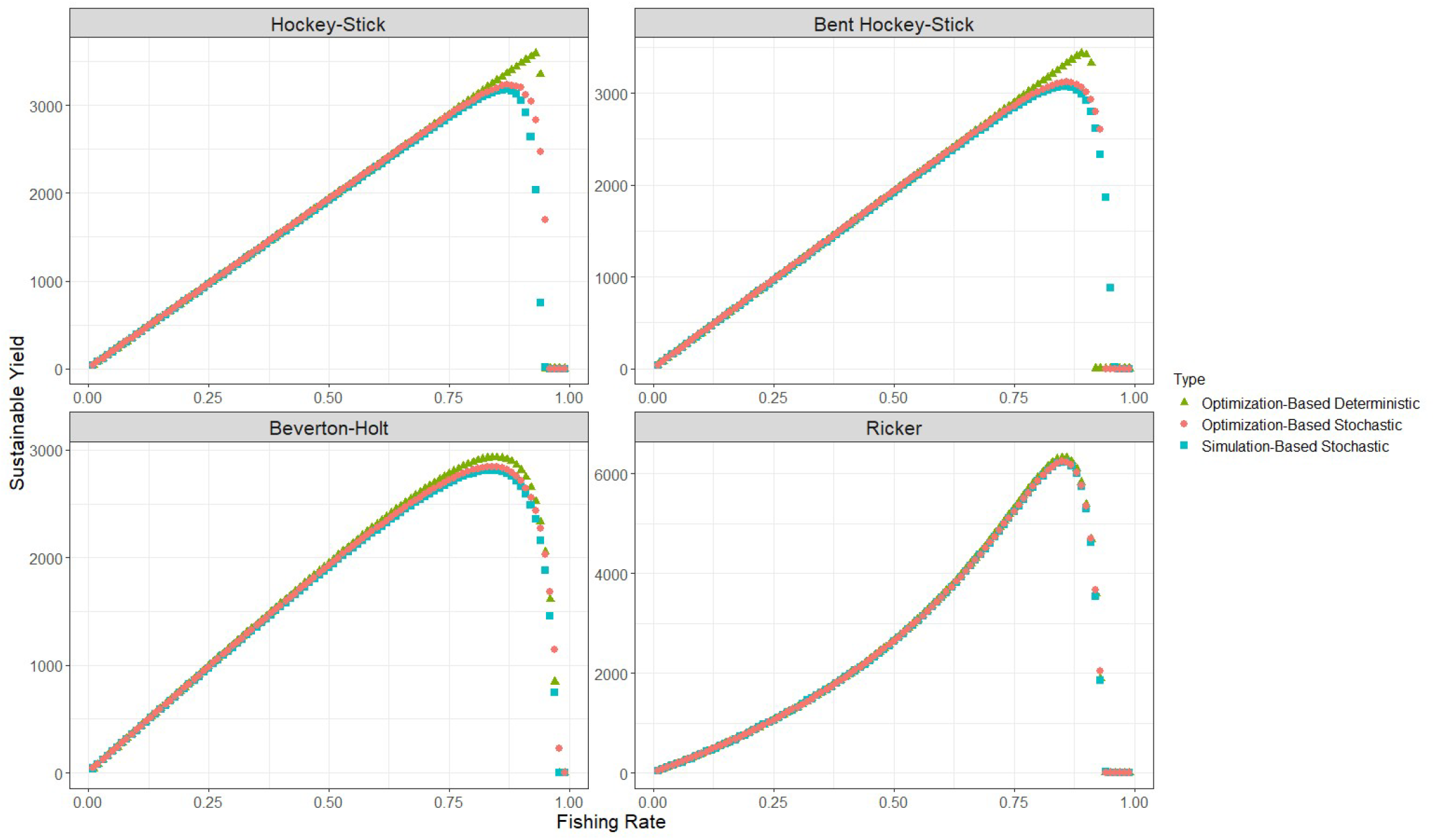
Surplus production curves against fishing rates obtained from stochastic simulation, optimization-based deterministic, and optimization-based stochastic calculations. Blue squares, green triangles, and red circles represent the simulation-based stochastic, optimization-based deterministic, optimization-based stochastic results, respectively.

In the simulation trials, the BHS model successfully converged with 100% probability under all settings, whereas the other SR functions (HS, MR, BH, and RI) failed to converge in some cases (the MR model failed in only one trial) (Table 2). The convergence failure of the HS model was likely due to non-differentiability at the breakpoint, indicating that the MR and BHS model properly circumvented this problem by smoothing the neighborhood around the breakpoint. When the BH model failed to converge, the SR curve became nearly flat. In contrast, when the RI model failed to converge, the gradients remained large, indicating that the parameter estimates were unstable.

The BHS model generally estimated biological reference points (*F*_*MSY*_ and *S*_*MSY*_) with small biases and standard deviations (Table 2; Figs. 4 and 5). The mean performance of the BHS model was the best among all models. The mean biases of HS, MR, BHS, BH, and RI across all trials were –2.39%, –1.90%, –0.61%, 2.24%, and -6.24% for *F*_*MSY*_, and 21.9%, 27.0%, 2.4%, –24.5%, and 108.0% for *S*_*MSY*_, respectively. The HS and MR models tended to underestimate *F*_*MSY*_ in the simulation where the HS or BHS model was the true model, whereas they overestimated *F*_*MSY*_ in the simulation where the RI model was the true model. The HS, MR, and BHS models tended to overestimate *S*_*MSY*_, except in the simulation where the RI model was the true model (Table 2; Figs. 4 and 5).

**Figure 4.**
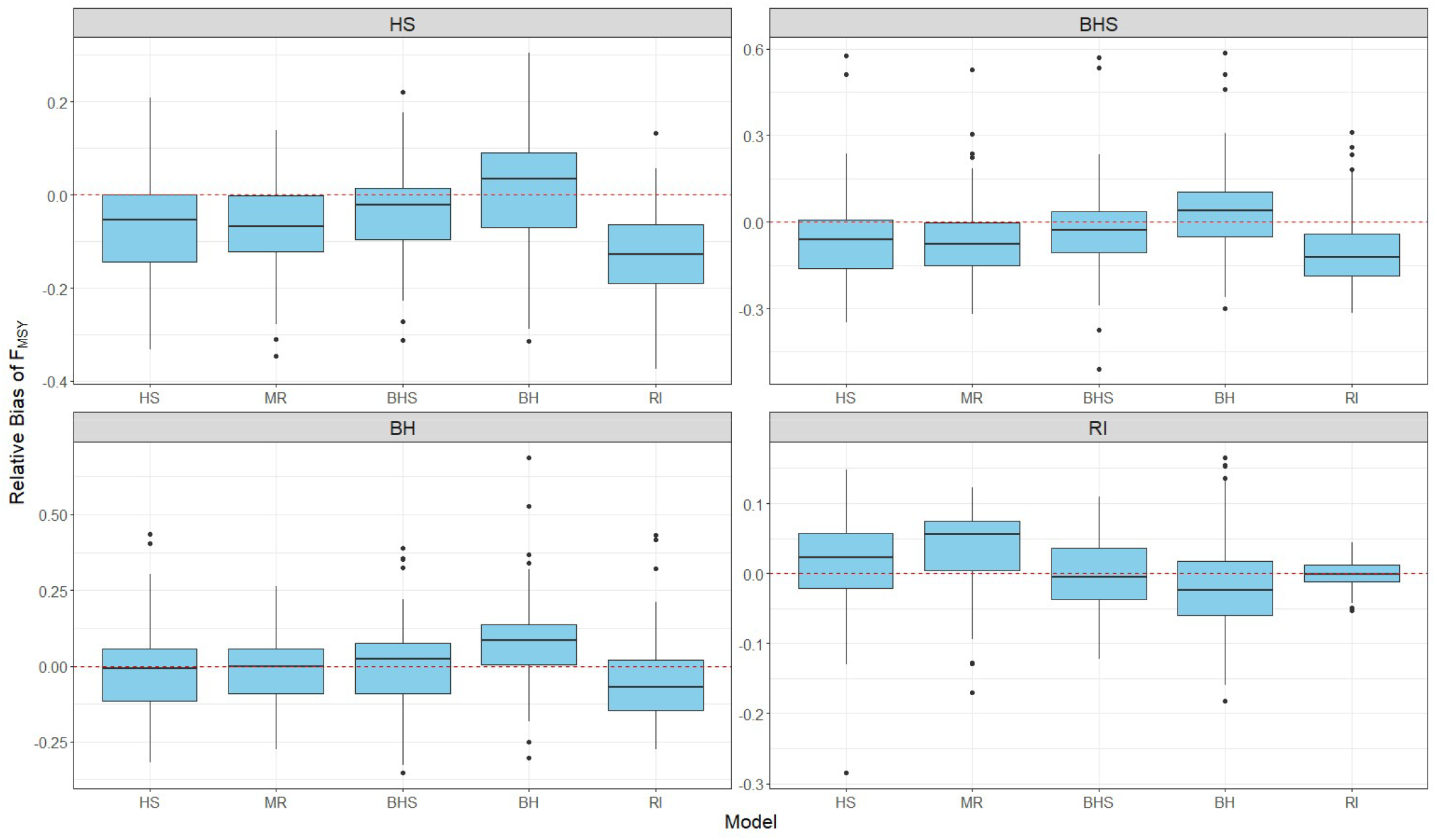
Relative bias of the fishing rate at MSY (*F*_*MSY*_) in the simulation. HS: Hockey-Stick; MR: Mesnil-Rochet; BHS: Bent Hockey-Stock; BH: Beverton-Holt: RI: Ricker. Each panel corresponds to the true SR function assumed in the simulation, and the x-axis represents the estimation model.

**Figure 5.**
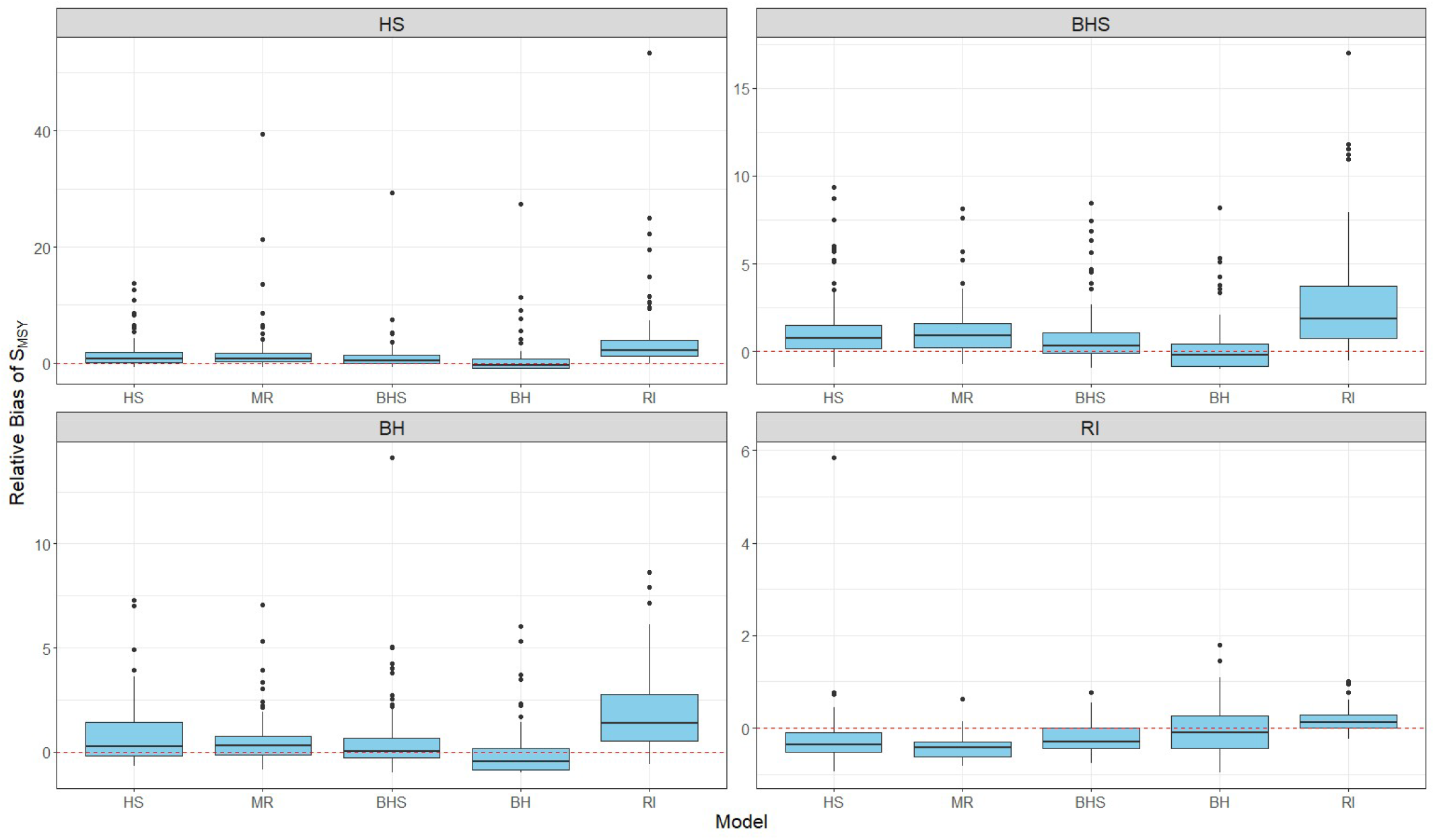
Relative bias of the spawning stock biomass at MSY in the simulation. Terminology and abbreviations are the same as in Figure 4.

The BH and RI models showed a clear contrast: the former generally overestimated *F*_*MSY*_ and underestimated *S*_*MSY*_, whereas the latter generally underestimated *F*_*MSY*_ and overestimated *S*_*MSY*_ (Table 2; Figs 4 and 5). In the simulation where the BH model was the true model, the fitted BH model underestimated the *b* parameter and overestimated the *a* parameter on average, indicating that density dependence was overestimated, i.e., estimated to be stronger than it actually was. In contrast, when the RI model was the true model, the fitted RI model slightly overestimated the *b* parameter on average, indicating that density dependence was underestimated, i.e., estimated to be weaker than it actually was.

Although the simulation results included some cases that failed to converge, excluding those cases did not substantially change the results; the figures obtained were similar to Figs. 4 and 5 (not shown here).

Although the *k*_*m*_ parameter was fixed at 5 in the present analysis, preliminary investigations suggest that this value is a reasonable choice for the available data (Supplementary Online Materials).

## Discussion

The BHS model successfully converged for all simulated datasets and generally showed better performance than the other models. The fitted BHS model even outperformed the HS and MR models when the HS model was the true model in the simulation, and it also outperformed them when the BHS model was the true model. Because the BHS model can flexibly adjust its curve shape by changing the *k*_*m*_ parameter while retaining the essential HS property —that recruitment becomes constant above the breakpoint— it is potentially more versatile than the traditional HS-type models. Moreover, the BHS model performed as well as or better than the BH and RI models even when those models were the true models, indicating its robustness to SR model misspecification (Figs. 4 and 5). The BHS model can take a functional form similar to that of the BH model to the left of the breakpoint by adjusting the *k*_*m*_ parameter, while also avoiding a completely flat or entirely linear (density-independent) curve.

The HS and MR models showed similar trends; *F*_*MSY*_ was underestimated and *S*_*MSY*_ was overestimated. Underestimation of maximum growth rates (the parameter *a*) estimated with the HS and MR models was likely to cause underestimation of *F*_*MSY*_ and overestimation of *S*_*MSY*_. Such biases are possibly due to scarce data below the breakpoint. On the other hand, the BH model overestimated *F*_*MSY*_ because it often overestimated maximum growth rates and predicted a completely flat curve. The general tendency for the BH model to overestimate *F*_*MSY*_ is consistent with the findings of Barrowman and Myers (2000). Because the BHS model can take an intermediate form between the HS/MR models and the BH model, it could provide an intermediate and nearly unbiased *F*_*MSY*_. This indicates that the BHS model would be more useful for highly productive species. While the RI model provided largely biased results when the other SR models were true, it was the best model when it was the true model. This implies that the RI model is sensitive to model misspecification.

We set the *k*_*m*_ parameter in the BHS model to 5 by default. When the BHS model failed to converge even after changing the initial parameter values, we varied the *k*_*m*_ parameter to achieve convergence. Because we used sparse and short time-series data as an example, it was difficult to achieve convergence across all possible ranges of the *k*_*m*_ parameter. Therefore, we determined the *k*_*m*_ parameter value arbitrarily in this study. Identifiability issues associated with the smoothness parameter *k*_*m*_ may affect both numerical convergence and the estimation of biological reference points, depending on data conditions. In short and sparse stock–recruitment time series, which are typical in many assessments, the likelihood surface with respect to *k*_*m*_ can be relatively flat, leading to weak identifiability. Under such conditions, estimation of *k*_*m*_ by maximum likelihood may result in convergence failures or unstable estimates, particularly for extremely small values of *k*_*m*_, as observed in our example (Supplementary Online Materials). Weak identifiability of *k*_*m*_ can also propagate into biological reference points. When *k*_*m*_ is poorly constrained, estimates of *F*_*MSY*_ and *S*_*MSY*_ may become sensitive to the assumed value of *k*_*m*_, potentially introducing bias if an inappropriate value is chosen. However, our sensitivity analysis in Supplementary Online Materials indicates that *F*_*MSY*_ and *S*_*MSY*_ were relatively stable for *k*_*m*_ values above a certain threshold, suggesting that the impact of identifiability on reference points may be limited when *k*_*m*_ is restricted to a reasonable range. As in nonparametric regression, smoothness or shape parameters are often treated as tuning parameters rather than parameters to be estimated by maximum likelihood (Green and Silverman, 1994). Accordingly, the *k*_*m*_ parameter should be treated as a tuning parameter, and an appropriate value of *k*_*m*_ could be determined using cross-validation (Green and Silverman, 1994) or time-series cross-validation (Hyndman and Athanasopoulos, 2019; Okamura et al., 2021). Practical guidance on the use of time-series cross-validation for selecting the *k*_*m*_ parameter is provided in the Supplementary Online Materials. With longer time series, broader coverage of spawning stock biomass, or lower process noise, identifiability of *k*_*m*_ is expected to improve, thereby reducing both convergence problems and the risk of bias in biological reference points.

MSY under stochastic environments (i.e., when population dynamics include multiplicative process noise) can differ from that under deterministic environments (i.e., without process noise) (Bousquet et al., 2008; Bordet and Rivest, 2014). Although analytical formulas are available for specific SR functions (Bordet and Rivest, 2014; Pedersen and Berg, 2017), they are not always attainable. The algorithm presented in this study is general and can be applied even to age-structured models. In particular, because biological reference points can vary substantially with the magnitude of process noise when hockey-stick-type SR functions are assumed, our new algorithm is especially useful in such cases.

A numerical algorithm for estimating biological reference points has several advantages, such as fast computation and independence from random seeds. Trijoulet et al. (2022) demonstrated that internal estimation of biological reference points within assessment models offers advantages over external estimation. However, because the per-recruit (PR) method used for internal estimation requires analytical formulas to calculate biological reference points (Albertsen and Trijoulet, 2020), our algorithm cannot be directly applied at present. To account for uncertainty in biological reference points, the delta method can be used, in which the approximate variance is obtained from the estimated variance-covariance matrix of the fixed parameters and the gradient vectors of biological reference points. Evaluating differences between internal and external approaches in terms of our estimation algorithm will be one of the future challenges.

Simulations based on the Falkland Islands’ *Loligo gahi* squid data suggested that using the BH model may be problematic from a conservation perspective, whereas using the RI model may be problematic from a fisheries efficiency perspective in managing this stock. For highly productive species such as squids, particularly when the available time series is short, using the BHS models can provide more stable estimates of biological reference points and offer a practical solution when the BH model yields an almost flat curve, regardless of the true SR relationship.

The HS-SR function is simple and has several advantages over traditional SR functions such as the BH and RI models. Its analytical drawback lies in the non-differentiability at the breakpoint. Although Mesnil and Rochet (2010) proposed a differentiable alternative, other functional forms can also reproduce characteristics similar to those of the original HS model. The model presented in this study incorporates a smoothed breakpoint, retaining the advantages of the HS model while providing greater robustness and flexibility than the Mesnil and Rochet’s bent-hyperbola SR function. Our BHS model is therefore promising as a new alternative. One caveat when using HS-type SR functions is that their biological reference points are highly sensitive to multiplicative process noise (Fig. 3). The new algorithm developed in this study for estimating biological reference points under stochastic environments is general and widely applicable to any SR model with multiplicative process noise. The combination of the new bent hockey-stick SR function and the new stochastic MSY estimation algorithm will greatly contribute to improving the robustness and sustainability of fisheries management.

## Acknowledgements

This work was supported by JSPS KAKENHI Grant Number JP 23K11025.

