## Supplement for "Bent Hockey-Stick Stock-Recruitment Function"

### 1. *Identifiability of the $k_m$ parameter*

We examined the identifiability of the $k_m$ parameter by evaluating the profile likelihood for the BHS model applied to the Falkland Islands' *Loligo gahi* squid stock from 1987 to 2000 (McAllister et al., 2004). A grid of $k_m$ values was considered: {0.1, 0.5, 1, 3, 5, 7, 10, 15, 20, 30}. The model failed to converge for $k_m = 0.1$ and 3. For the remaining values, the negative log-likelihood was largely flat over a wide range of $k_m$ values (approximately 1–15; Fig. S1). This flatness indicates that the $k_m$ parameter is only weakly identifiable and cannot be reliably estimated using a maximum likelihood approach with these data. Consequently, this result motivates treating $k_m$ as a tuning parameter rather than estimating it directly by maximum likelihood.

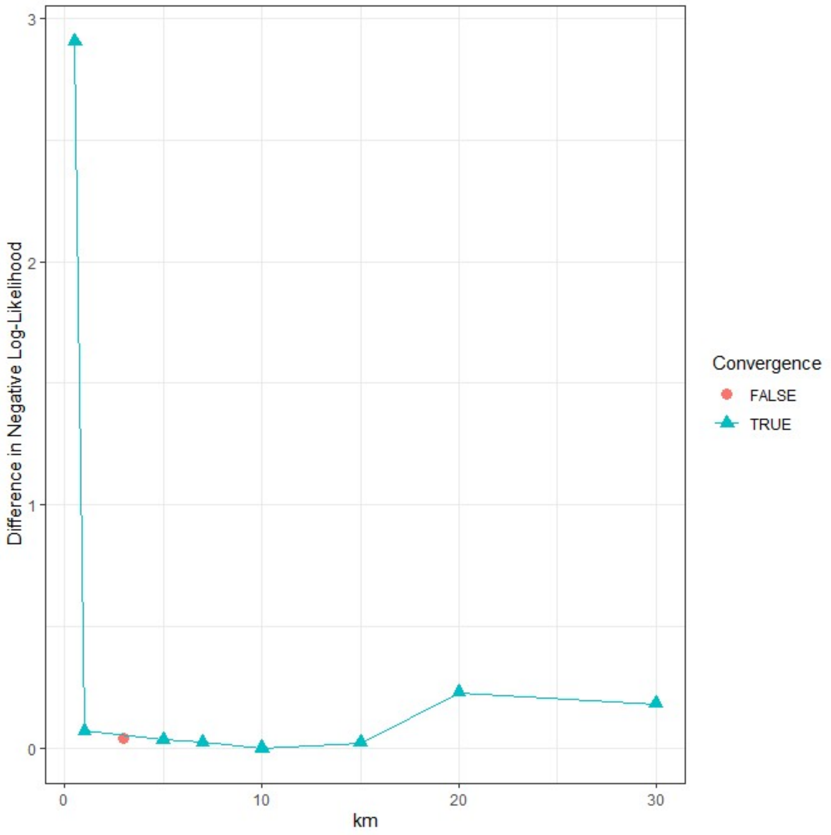


Fig. S1. Negative (marginal) log-likelihood as a function of the $k_m$ value. The model failed to converge for $k_m = 0.1$ and 3. The result for $k_m = 0.1$ (final objective function $\approx 177$) was omitted for clarity.

### 2. *Sensitivity of stock–recruitment curves and biological reference points to the $k_m$ parameter*

The stock–recruitment (SR) curves corresponding to different values of $k_m$ are shown in Fig. S2. For $k_m$ = 0.5, the estimated asymptotic recruitment was substantially large; however, this result was associated with a relatively low likelihood, suggesting that the estimate is poorly supported by the data. The SR curves for $k_m$ values between 1 and 15 were broadly similar, whereas those for $k_m$ greater than 15 differed more markedly, consistent with the differences in log-likelihood values (Fig. S1).

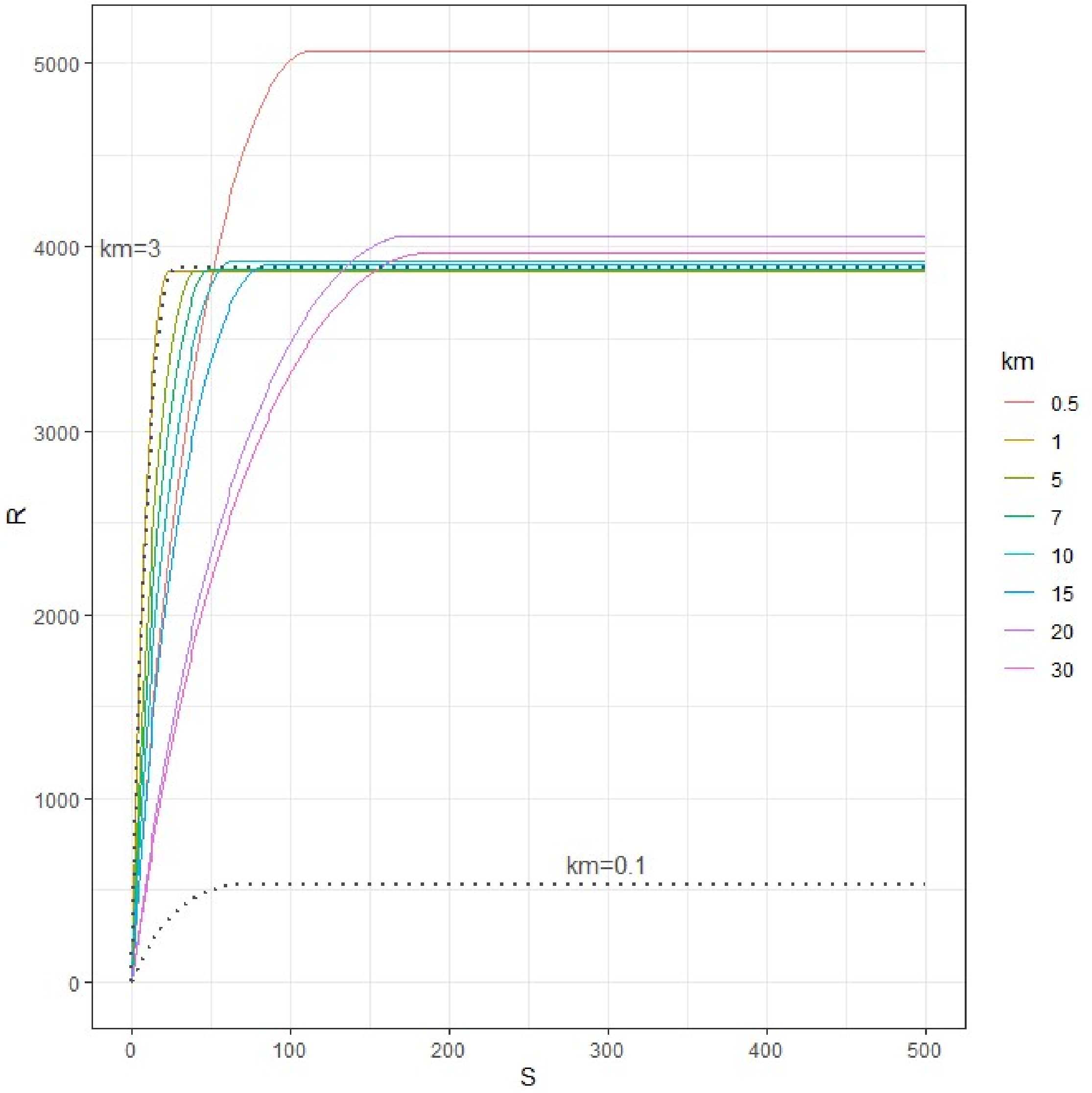


Fig. S2. Stock-recruitment curves for different values of $k_m$. Gray dotted curves correspond to results for non-converged $k_m$ values (0.1 and 3).

We calculated $F_{MSY}$ and $S_{MSY}$ as biological reference points for different values of $k_m$ (Figs. S3 and S4). Both $F_{MSY}$ and $S_{MSY}$ were stable for $k_m$ values equal to or greater than 5. This suggests that choosing $k_m$ values of 5 or higher leads to stable estimation of biological reference points for these data.

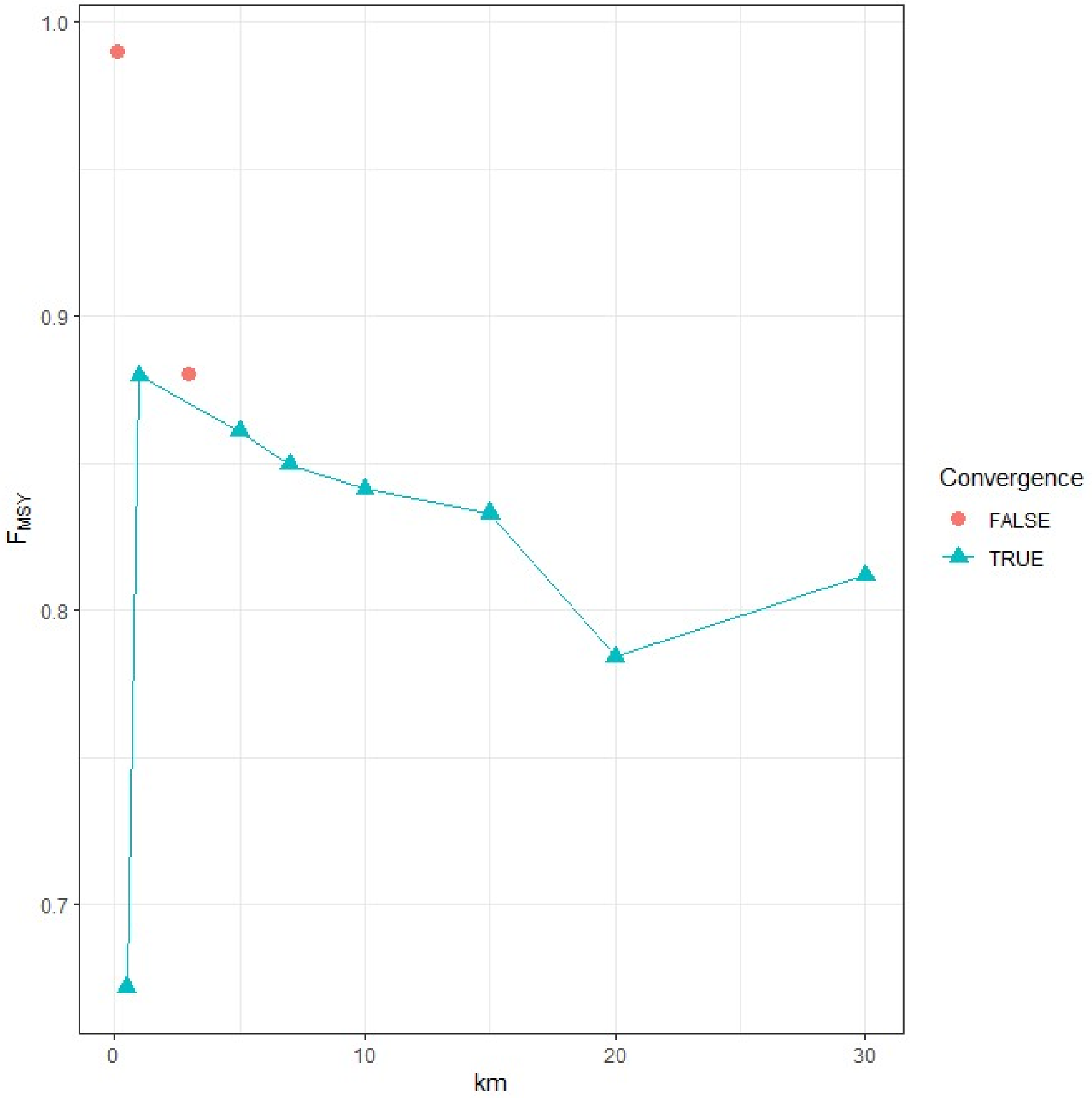


Fig. S3. $F_{MSY}$ for different values of $k_m$.

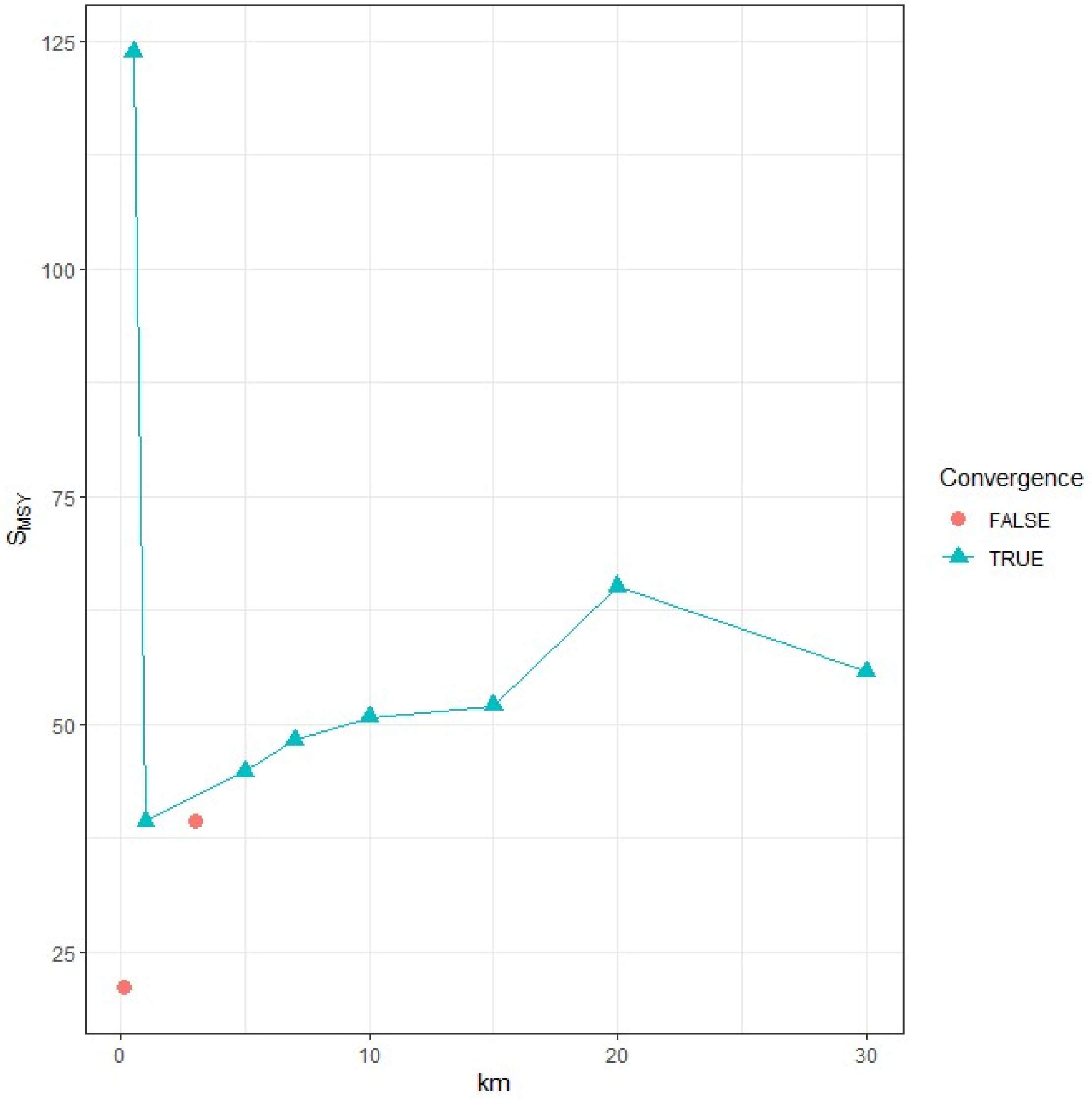


Fig. S4. $S_{MSY}$ for different values of $k_m$.

#### 3. *Preliminary analysis for selecting an appropriate value of the $k_m$ parameter*

Shape parameters are generally difficult to estimate reliably using a maximum likelihood approach, particularly when the likelihood surface is relatively flat. We therefore applied time-series cross-validation (Hyndman and Athanasopoulos 2019), also referred to as retrospective forecasting (RF; Brooks and Legault 2016), to explore an appropriate value of $k_m$. The $k_m$ parameter was evaluated by visually inspecting the following retrospective forecasting error metric (Okamura et al., 2021):

$$RF_R = \exp\left(\frac{1}{P}\sum_{t=1}^{P} \log\left[\left(\hat{r}_{T-(t-1)}^{1:T} - \hat{r}_{T-(t-1)}^{1:(T-t)}\right)^2\right]\right),$$

where $T$ is the length of the time series, $\hat{r}^{1:T}_{T-(t-1)}$ is the predicted log-recruitment in year $T-(t-1)$ from the analysis result using the full dataset, and $\hat{r}^{1:(T-t)}_{T-(t-1)}$ is the corresponding prediction obtained using data from years 1 to $T-t$. $P$ denotes the number of retrospective steps. We provisionally set $P = 4$ because of the short time series ($T = 14$).

The optimal $k_m$ value was 5 among the candidate values. In contrast to the profile likelihood, the RF errors showed clearer differences among $k_m$ values, suggesting that RF provides more discriminative information for tuning the smoothness parameter with short time-series data. Although this analysis is preliminary and based on a single dataset, the clearer separation observed in RF errors compared with the profile likelihood suggests that retrospective forecasting may provide a practical and broadly useful approach for tuning $k_m$ when likelihood-based identifiability is weak.

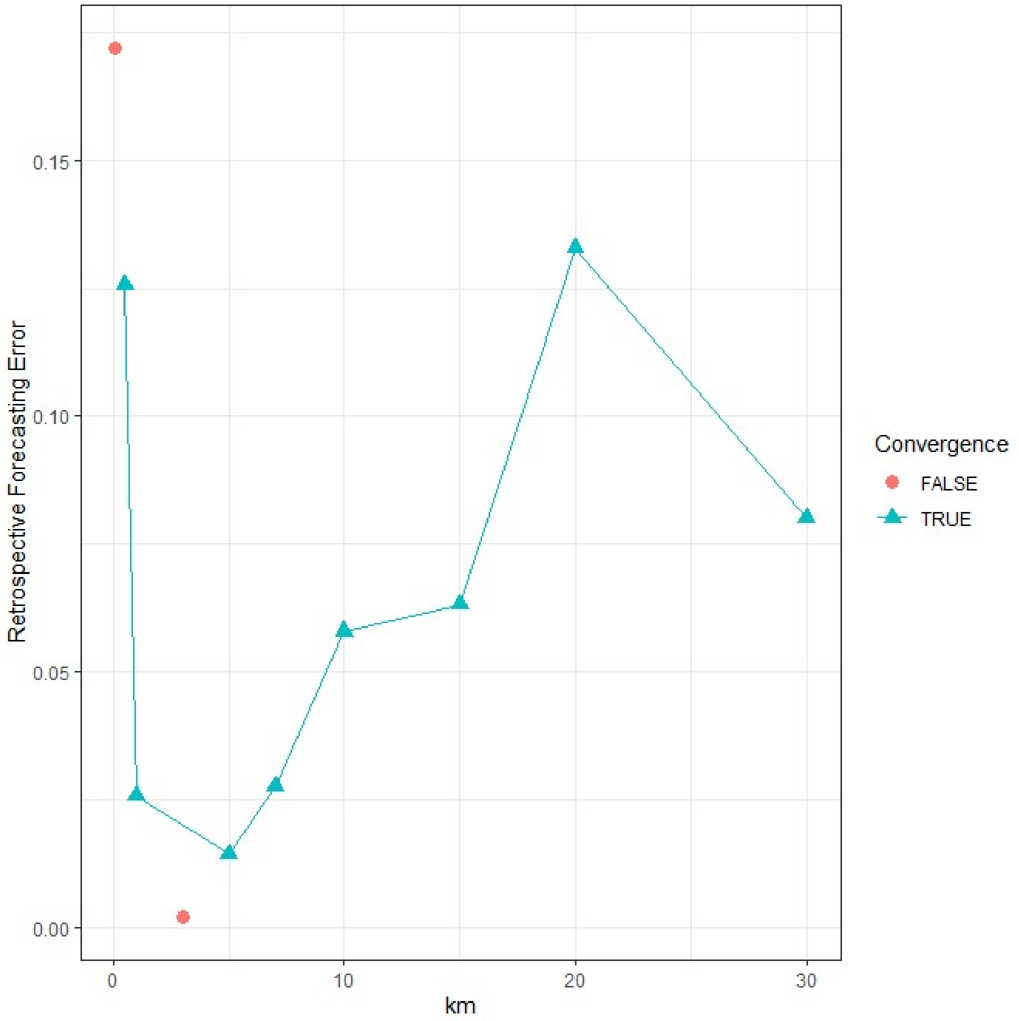


Fig. S5. Retrospective forecasting errors for different values of $k_m$.
